# Comparison of two ancient DNA extraction protocols for skeletal remains from tropical environments

**DOI:** 10.1101/184119

**Authors:** Maria A. Nieves-Colón, Andrew T. Ozga, William J. Pestle, Andrea Cucina, Vera Tiesler, Travis W. Stanton, Anne C. Stone

## Abstract

**Objectives:** The tropics harbor a large part of the world’s biodiversity and have a long history of human habitation. However, paleogenomics research in these climates has been constrained so far by poor ancient DNA yields. Here we compare the performance of two DNA extraction methods on ancient samples of teeth and petrous portions excavated from tropical and semitropical sites in Tanzania, Mexico, and Puerto Rico (N=12).

**Materials and Methods:** All samples were extracted twice, built into double-stranded sequencing libraries, and shotgun sequenced on the Illumina HiSeq 2500. The first extraction protocol, Method D, was previously designed for recovery of ultrashort DNA fragments from skeletal remains. The second, Method H, modifies the first by adding an initial EDTA wash and an extended digestion and decalcification step.

**Results:** No significant difference was found in overall ancient DNA yields or post-mortem damage patterns recovered from samples extracted with either method, irrespective of tissue type. However, Method H samples had higher endogenous content and more mapped reads after quality-filtering, but also higher clonality. In contrast, samples extracted with Method D had shorter average DNA fragments.

**Discussion:** Both methods successfully recovered endogenous ancient DNA. But, since surviving DNA in ancient or historic remains from tropical contexts is extremely fragmented, our results suggest that Method D is the optimal choice for working with samples from warm and humid environments. Additional optimization of extraction conditions and further testing of Method H with different types of samples may allow for improvement of this protocol in the future.

---

Ancient DNA (aDNA) is a low quality and low quantity source of genetic material, which is highly susceptible to external contamination (Gilbert et al., 2006; Hofreiter et al., 2001; Pääbo, 1989). Due to the variety of taphonomic and diagenetic processes that take place after death, DNA decays exponentially once cell repair functions cease in biological tissues (Hofreiter et al., 2001). Consequently, most genetic information obtained from ancient samples is contained in small, degraded DNA fragments (Allentoft et al., 2012; Briggs et al., 2007; Dabney et al., 2013a). Because of this, until recently, aDNA studies have focused on short but informative fragments of the autosomal genome or, alternatively, on multicopy loci such as mitochondrial DNA (Ho & Gilbert, 2010). Recent advances in DNA extraction, target enrichment, and next-generation sequencing methods now allow the recovery of complete genomes from remains dating as far back in time as the early Holocene and Middle Pleistocene (Meyer et al., 2016; Meyer et al., 2014; Orlando et al., 2013).

Despite these improvements in stretching the time depth for aDNA recovery, paleogenomics research continues to be constrained in its geographic focus because DNA preservation is negatively correlated with thermal age due to the accelerating effect of high temperatures on biomolecule decay and fragmentation (Adler et al., 2011; Allentoft et al., 2012; Hofreiter et al., 2015; Kistler et al., 2017; Lindahl, 1993; Smith et al., 2001). Therefore, most aDNA studies focus on archaeological remains excavated from cold and temperate world regions, which have the highest chance of DNA survival (Paijmans et al., 2013; Wade, 2015). Nevertheless, despite the challenges of working with poorly preserved samples, several studies have successfully recovered aDNA from tropical sites in the Caribbean, the Yucatan peninsula and South-East Asia (Damgaard et al., 2015; Gamba et al., 2014; Gutierrez-Garcia et al., 2014; Kehlmaier et al., 2017; Mendisco et al., 2015; Schroeder et al., 2015). Both today and in the past, the tropics harbor a large portion of the world’s biodiversity and human settlements (Brown, 2014; Buzas et al., 2002). Therefore, understanding how DNA is preserved in degraded remains excavated from these environments and optimizing or improving methods that facilitate aDNA recovery from these contexts is of great interest to advance anthropology, paleontology, and conservation genetics, among other fields.

Here we test the performance of two DNA extraction protocols on tooth and petrous portion samples from degraded skeletal remains recovered from tropical sites in Tanzania, Mexico and Puerto Rico (N=12). Specifically, we compare the method developed by (Dabney et al., 2013a), hereafter Method D, to a second approach, Method H. Method H modifies the former by adding an initial EDTA wash, as in Warinner et al. (2014), and an extended digestion and decalcification step as in Gamba et al. (2016). Method D was specifically designed to increase recovery of extremely short DNA fragments (as small as 30 base pairs) in ancient bone and tooth extractions. In line with earlier protocols (Höss & Pääbo, 1993; Rohland & Hofreiter, 2007), this method employs a 24-hour proteinase K digestion to break up cell proteins, and uses a chaotropic guanidium-based salt to bind DNA fragments and remove inhibitors. It differs from previous approaches in its use of silica spin columns and a guanidine hydrochloride binding buffer (instead of guanidine thiocyanate). Method D has been successfully employed in the recovery of aDNA from Late Pleistocene cave bear remains (Dabney et al., 2013a), from Middle Pleistocene hominin fossils (Meyer et al., 2016; Meyer et al., 2014), and from a large variety of more recently dated human and animal remains, including at least one from a tropical context (Günther et al., 2015; Heintzman et al., 2015; Kehlmaier et al., 2017; Seguin-Orlando et al., 2014).

Since Method D was developed, several extraction protocol modifications have been proposed for improving endogenous aDNA recovery. Comparing different extraction methods, Gamba et al. (2016) found that a secondary digestion and decalcification using lysis buffer with EDTA, proteinase K and N-laurylsarcosyl detergent solution, aided in solubilizing cell proteins and resulted in increased aDNA yields. Similarly, Damgaard et al. (2015) observed that a brief pre-digestion (between 15 and 30 minutes) with an EDTA and proteinase K buffer was successful in reducing proportions of exogenous, contaminant DNA and in enriching extracts for endogenous aDNA. The use of similar detergent solutions has been implemented previously in extraction protocols designed by Richards et al. (1995) and was also recently reported in extractions of petrous portion tissue (Gamba et al., 2014; Pinhasi et al., 2015). Likewise, Warinner et al. (2014) used an initial EDTA wash to remove loosely bound surface contaminants on mineralized dental calculus without significant DNA loss. This finding was mirrored by Tromp et al. (2017) who observed that EDTA decalcification was more effective at recovering microparticles from dental calculus than hydrochloric acid.

In this study, we evaluate whether using a modified version of the Method D protocol (henceforth Method H), with an initial EDTA wash and an extended digestion and decalcification step, results in improved endogenous aDNA recovery. In addition, because the majority of development in aDNA extraction protocols has been conducted with samples from temperate or cold contexts (Barlow et al., 2016; Boessenkool et al., 2016; Dabney et al., 2013a; Gamba et al., 2016; Gamba et al., 2014; Glocke & Meyer, 2017) but see (Damgaard et al., 2015; Pinhasi et al., 2015; Tromp et al., 2017), this work focuses on protocol optimization with tooth and petrous portion samples recovered from tropical sites. Here we examine raw DNA yields and endogenous reads, recovered after shotgun Illumina sequencing from parallel, paired extractions and characterize differences in base pair composition, sequence read complexity, post-mortem damage profiles, and average read lengths recovered between the two methods. Archaeological samples included in this research were obtained from human remains excavated at three Ceramic Age sites from Puerto Rico (n=5) and one tomb from the Maya site of Yaxuna in Yucatán, Mexico (n=6). Additionally, one historic sample from a Tanzanian chimpanzee was also included.

Study results suggest that both methods were similarly efficient at aDNA recovery. Libraries sequenced from Method H extracts had higher endogenous content but also higher clonality, while libraries sequenced from Method D extracts have shorter DNA fragments. Since most of the archaeological samples had extremely low endogenous content (<5%), and average DNA fragment sizes were under 80 bp, we conclude that Method D is better suited than Method H for maximized recovery of informative ancient DNA molecules from remains buried in tropical environments. However, one important caveat of our study is that these findings are only applicable to tooth samples, since the small sample of petrous portions obtained led to inconclusive results in statistical tests conducted with this tissue.

## MATERIALS AND METHODS

### Sample and site information

Samples were collected from skeletal remains excavated from three tropical contexts. Tissue samples from petrous portions and/or teeth were obtained from six human skeletons excavated from a single tomb in the archaeological site of Yaxuna in Yucatán, Mexico. This was an originally unfilled burial space that dates to the Maya Early Classic period (specifically from the 6^th^ century A.D.) During the centuries of deposition, the burial space gradually filled with rubble and fill falling from the ceiling of the chamber. In most skeletons, only one tissue type, petrous portion or teeth was available for sampling, while only two individuals (AD-372 and AD-373) could be sampled in both anatomic locations (Table 1). Additionally, five teeth were collected from humans remains excavated from three open-air sites in Puerto Rico: Tibes (n=1), Paso del Indio (n=2), and Punta Candelero (n=2). All five individuals date from pre-contact Ceramic Age contexts, between A.D. 500-1300 (Pestle, 2010; Pestle & Colvard, 2012). Lastly, one tooth was collected from the skeletal remains of a wild chimpanzee (*Pan troglodytes schweinfurthii*) who died in 1966. The chimpanzee was named McDee by scientific observers in western Tanzania (prior to Gombe becoming a national park) and is referred to in this study as GB-7. After death, MacDee’s remains were stored in a metal box. Flesh was eaten away by insects and the skeleton was subsequently cleaned. Petrous portion tissue was not available from the Puerto Rican or Tanzanian remains. In total, 12 individual skeletons were sampled producing ten teeth and four petrous portions.

**Table 1.**
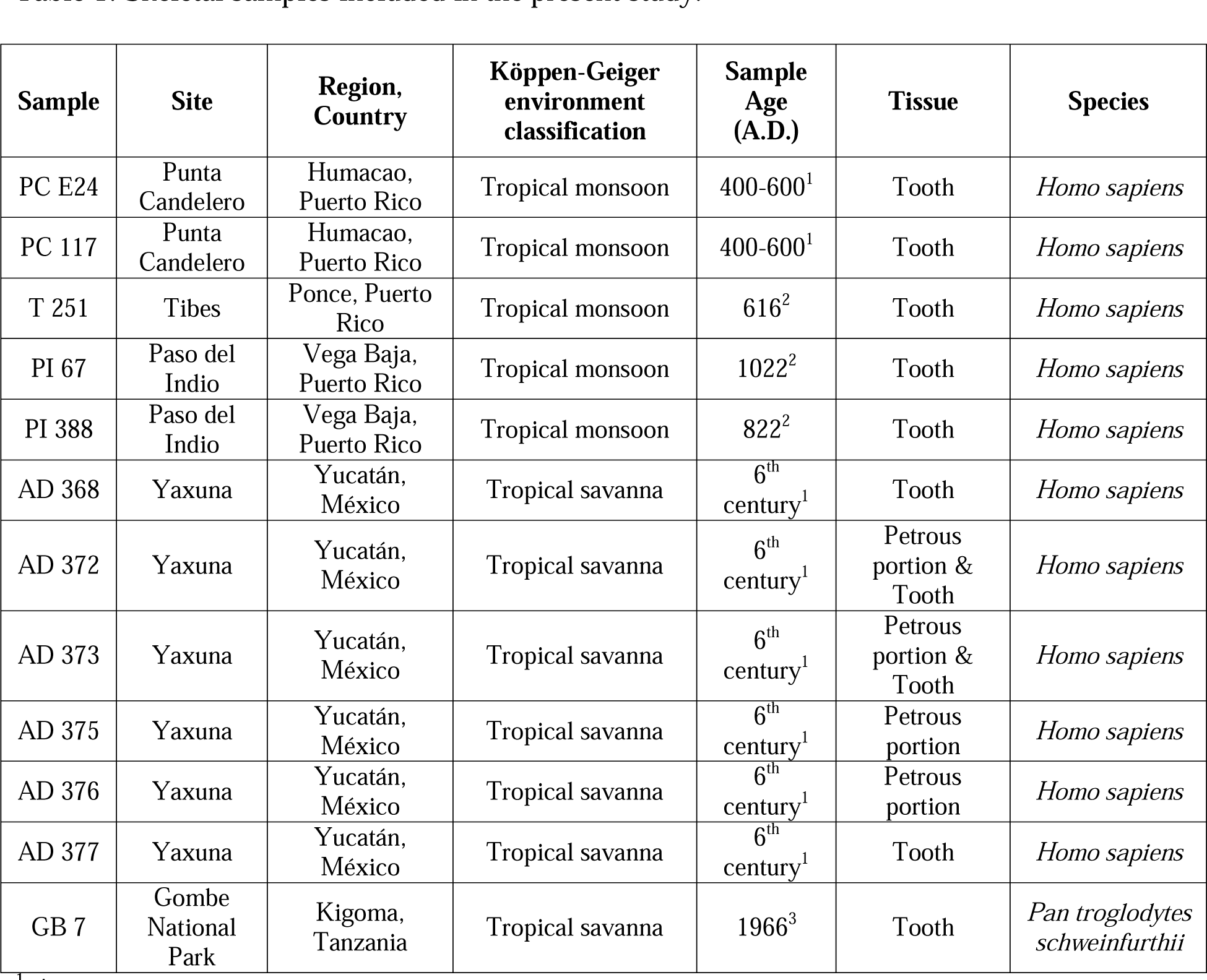
Skeletal samples included in the present study.

| Sample | Site | Region,<br>Country | Köppen-Geiger<br>environment<br>classification | Sample<br>Age<br>(A.D.) | Tissue | Species |
| --- | --- | --- | --- | --- | --- | --- |
| PC E24 | Punta<br>Candelero | Humacao,<br>Puerto Rico | Tropical monsoon | 400-600<sup>1</sup> | Tooth | *Homo sapiens* |
| PC 117 | Punta<br>Candelero | Humacao,<br>Puerto Rico | Tropical monsoon | 400-600<sup>1</sup> | Tooth | *Homo sapiens* |
| T 251 | Tibes | Ponce, Puerto<br>Rico | Tropical monsoon | 616<sup>2</sup> | Tooth | *Homo sapiens* |
| PI 67 | Paso del<br>Indio | Vega Baja,<br>Puerto Rico | Tropical monsoon | 1022<sup>2</sup> | Tooth | *Homo sapiens* |
| PI 388 | Paso del<br>Indio | Vega Baja,<br>Puerto Rico | Tropical monsoon | 822<sup>2</sup> | Tooth | *Homo sapiens* |
| AD 368 | Yaxuna | Yucatán,<br>México | Tropical savanna | 6<sup>th</sup><br>century<sup>1</sup> | Tooth | *Homo sapiens* |
| AD 372 | Yaxuna | Yucatán,<br>México | Tropical savanna | 6<sup>th</sup><br>century<sup>1</sup> | Petrous<br>portion &<br>Tooth | *Homo sapiens* |
| AD 373 | Yaxuna | Yucatán,<br>México | Tropical savanna | 6<sup>th</sup><br>century<sup>1</sup> | Petrous<br>portion &<br>Tooth | *Homo sapiens* |
| AD 375 | Yaxuna | Yucatán,<br>México | Tropical savanna | 6<sup>th</sup><br>century<sup>1</sup> | Petrous<br>portion | *Homo sapiens* |
| AD 376 | Yaxuna | Yucatán,<br>México | Tropical savanna | 6<sup>th</sup><br>century<sup>1</sup> | Petrous<br>portion | *Homo sapiens* |
| AD 377 | Yaxuna | Yucatán,<br>México | Tropical savanna | 6<sup>th</sup><br>century<sup>1</sup> | Tooth | *Homo sapiens* |
| GB 7 | Gombe<br>National<br>Park | Kigoma,<br>Tanzania | Tropical savanna | 1966<sup>3</sup> | Tooth | *Pan troglodytes*<br>*schweinfurthii* |

<sup>1</sup> Approximate date, based on archaeological context.
<sup>2</sup> Radiocarbon date median probability calAD.
<sup>3</sup> Date of individual death.

^1^ Approximate date, based on archaeological context.
^2^ Radiocarbon date median probability calAD.
^3^ Date of individual death.

### Sample processing and DNA extraction

Sample processing and DNA extractions were conducted at the Arizona State University Ancient DNA Laboratory, a Class 10,000 clean-room facility. To eliminate surface contaminants and inhibitors, tooth and bone samples were cleaned with a 1% sodium hypochlorite solution. The outer surface was mechanically removed with a Dremel tool (Rohland & Hofreiter, 2007) and samples were UV irradiated for 5 minutes on each side in a UVP CL-1000 Ultraviolet Crosslinker. Teeth were sliced transversally at the cemento-enamel junction using a Dremel tool. The roots were covered in aluminum foil and pulverized by blunt force with a hammer as in Schuenemann et al. (2011). Petrous portions were sampled as recommended by Pinhasi et al. (2015). All laboratory procedures were conducted using contamination controls, such as use of full body coverings, bleach decontamination and UV irradiation of tools and work area before and between uses (Cooper & Poinar, 2000; Gilbert et al., 2006).

Each sample was extracted twice, one time using Method D and a second time using Method H. Method D was implemented as in (Dabney et al., 2013a) with the modification that the TET buffer was warmed to 65 °C in a heat block. Method H combines steps from earlier protocols including an initial EDTA wash as in Warinner et al. (2014), an extended digestion and decalcification step as in Gamba et al. (2016), and binding and purification steps as in (Dabney et al., 2013a). See Supplementary File S1 for complete Method H protocol. Approximately 100 mg of bone or tooth powder were used for each extraction. 1 μl of each extract was used to measure DNA yields through fluorometric quantification with the Qubit 2.0 High Sensitivity assay (Table S1) (Simbolo et al., 2013). Extraction blanks were included throughout the process to monitor potential contamination.

### Library preparation and sequencing

Double stranded libraries were built following the protocol by Meyer and Kircher (2010) and including negative controls. DNA content in the libraries was quantified using real-time PCR (qPCR) with the 2X Dynamo SYBR Green qPCR Master Mix to determine the ideal amount of indexing cycles. All libraries were double indexed and amplified for 11-25 cycles following published guidelines (Kircher et al., 2012; Seguin-Orlando et al., 2015). Indexed libraries were purified using the Qiagen MinElute PCR purification kit, and DNA content after amplification was determined via qPCR using the KAPA Library Quantification kit following manufacturer instructions (Kapa Biosystems). Fragment analysis of the indexed libraries was conducted with the DNA 1000 assay on the Agilent 2100 Bioanalyzer. Heteroduplexes that arose during indexing were eliminated through reconditioning PCR and all libraries were purified and requantified as detailed above. Reconditioned shotgun libraries were sequenced on one lane across two runs on the Illumina HiSeq 2500 in Rapid Run mode (2 × 100 bp reads) at the Yale Center for Genomic Analysis. See Supplementary File S1 for additional details of library preparation, PCR primers and conditions.

### Shotgun read mapping and processing

Illumina sequence reads were merged and adapters trimmed using SeqPrep (https://github.com/istiohn/SeqPrep). To compare sequencing results directly across extraction treatments and to control for differences in sequencer output, 1 million reads were randomly selected for all samples using seqtk with default parameters (https://github.com/lh3/seqtk). For the human samples, the downsampled reads were mapped to the GRCh37 (hg19) reference with the mitochondria replaced by the revised Cambridge Reference Sequence (rCRS) (Andrews et al., 1999). For the chimpanzee samples, the reads were mapped to the PanTro4 assembly. Mapping was performed using BWA v. 0.7.5 (Heng Li & Durbin, 2009) following recommendations by Schubert et al. (2014). Duplicate reads were identified using the MarkDuplicates module v.2.12.1 within Picard Tools (http://broadinstitute.github.io/picard). Quality filtering (≥ Q30), removal of duplicates and of reads with multiple mappings was performed with SAMtools v. 0.1.19 (H. Li et al., 2009). BAM files were rescaled and damage patterns were characterized using mapDamage v.2.0.2 (Ginolhac et al., 2011; Jónsson et al., 2013). Parameters examined included deamination patterns, probability of C to T misincorporations at first position, probability of G to A misincorporations at last position, probability of a DNA fragment terminating in a single-stranded overhang (λ), probability of observing cytosine deamination in a double strand (δ_D_), and probability of observing cytosine deamination in a single strand context (δ_S_). Library complexity estimates were generated using *preseq* v2.0 (Daley & Smith, 2013) on downsampled BAM files containing all mapped reads. Summary statistics were estimated on rescaled BAM files using Qualimap v.2.2.1 (Okonechnikov et al., 2016). See Supplementary File S2 for additional details of shotgun sequence read processing.

Future experiment yield predictions were calculated for the two libraries with highest number of reads: GB-7 and PI-67. For these two samples, random downsampling was repeated matching the lowest number of reads obtained per sample-treatment combination after adapter trimming and merging: GB-7: 5,192,848 reads and PI-67: 6,876,556 reads. Read mapping and quality filtering were repeated for these data using the same parameters listed above. Extrapolation curves were calculated in *preseq* using the default step size parameter (−s 1000000) and extrapolating to 600,000,000 total reads. This is the maximum number of paired-end reads produced on a single flow-cell of the Illumina HiSeq 2500 in Rapid Run mode. For all other paired sample-treatment combinations extrapolation estimates failed due to low read depth.

### Statistical Analyses

Extraction yields (ng/μL), number of mapped, unique reads (defined as a mapped sequence read that has unique external coordinates), percent endogenous content, library complexity (percent distinct reads as measured by the *preseq* ccurve function), clonality (measured as fraction of the mapped reads that are duplicates: number of duplicate reads / number of mapped reads), percent GC content, average fragment lengths, and damage parameters were compared for each sample across extraction treatments by using paired T tests or non-parametric Wilcoxon signed rank tests. For these analyses, samples were subdivided according to type of tissue. Normality assumptions were evaluated using a Shapiro-Wilk normality test (Table S3) and through visual examination of Quantile-Quantile plots and histograms of the difference between paired values as recommended by Ghasemi and Zahediasl (2012) (Figure S1 and S2). Correlations between variables were tested using Pearson’s *r* as implemented in the R cor.test function.

### Computational resources and R packages

This research was conducted using resources from the ASU High Performance Computing Saguaro environment. All calculations were performed in R 3.2.4. Scripts written for this project are available at: https://github.com/mnievesc/aDNAExtMethodsPaper_scripts. All plots and figures were generated using the ggplot2 (H Wickham, 2009), gridExtra (Auguie, 2016), tidyr (H Wickham, 2016) and reshape 2 (Hadley Wickham, 2007) packages or with R base graphics (R Core Team, 2016).

## RESULTS

### DNA yields

DNA yields were evaluated through flourometric quantifications of raw extracts (ng/μL) (Figure S3). These analyses did not reveal significant differences in mean DNA yields between samples extracted with either method irrespective of tissue (Table 2).

**Table 2.** Results of paired tests. Significant values P<0.05 bolded.

| Test comparison | Tissue | Paired test | Test statistic | DF | P-value |
| --- | --- | --- | --- | --- | --- |
| DNA yields (ng/μL) | Tooth | T-test | $t = -0.1924$ | 9 | 0.8517 |
| | Petrous portion | T-test | $t = -1.2219$ | 3 | 0.3090 |
| Number of mapped, unique reads | Tooth | Wilcoxon signed ranks | $V = 4$ | n/a | **0.0136** |
| | Petrous portion | Wilcoxon signed ranks | $V = 0$ | n/a | 0.1250 |
| Percent endogenous content | Tooth | Wilcoxon signed ranks | $V = 4.5$ | n/a | **0.0217** |
| | Petrous portion | Wilcoxon signed ranks | $V = 0$ | n/a | 0.1250 |
| Percent clonality | Tooth | Wilcoxon signed ranks | $V = 7$ | n/a | **0.0371** |
| | Petrous portion | T-test | $t = -0.3852$ | 3 | 0.7257 |
| Percent distinct reads | Tooth | Wilcoxon signed ranks | $V = 36$ | n/a | 0.1232 |
| | Petrous portion | T-test | $t = 0.5300$ | 3 | 0.6328 |
| Average fragment length | Tooth | Wilcoxon signed ranks | $V = 2$ | n/a | **0.0058** |
| | Petrous portion | Wilcoxon signed ranks | $V = 1$ | n/a | 0.2500 |
| Percent average GC content | Tooth | Wilcoxon signed ranks | $V = 46$ | n/a | 0.0644 |
| | Petrous portion | Wilcoxon signed ranks | $V = 10$ | n/a | 0.1250 |
| Damage parameter $\delta_D$ | Tooth | Wilcoxon signed ranks | $V = 30$ | n/a | 0.8457 |
| | Petrous portion | Wilcoxon signed ranks | $V = 3$ | n/a | 0.625 |
| Damage parameter $\delta_S$ | Tooth | Wilcoxon signed ranks | $V = 35$ | n/a | 0.4922 |
| | Petrous portion | Wilcoxon signed ranks | $V = 6$ | n/a | 0.8750 |
| Damage parameter $\lambda$ | Tooth | Wilcoxon signed ranks | $V = 42$ | n/a | 0.1601 |
| | Petrous portion | Wilcoxon signed ranks | $V = 4$ | n/a | 0.8750 |

### Endogenous content and library complexity

For each DNA library, between 1 and 27 million reads were obtained after shotgun sequencing. For consistency, statistical analyses were performed with a starting number of one million randomly selected reads for all samples. Percentage endogenous content was calculated as the proportion of unique reads mapping to the reference (after duplicate removal and quality filtering) over the total amount of down sampled reads (Table S2). Most samples had <5% endogenous content, except for the chimpanzee sample, GB-7, which yielded >10% endogenous content, an up to sixteen-fold higher content than that found in the human libraries. This difference may be attributable to the younger age of the historic chimpanzee sample. Overall, samples extracted with Method H tended to have higher endogenous content and more unique reads mapping to the reference post-quality filtering (Figure 1, Figure S4). These differences were statistically significant for teeth but not petrous portion samples (Table 1).

**Figure 1.**
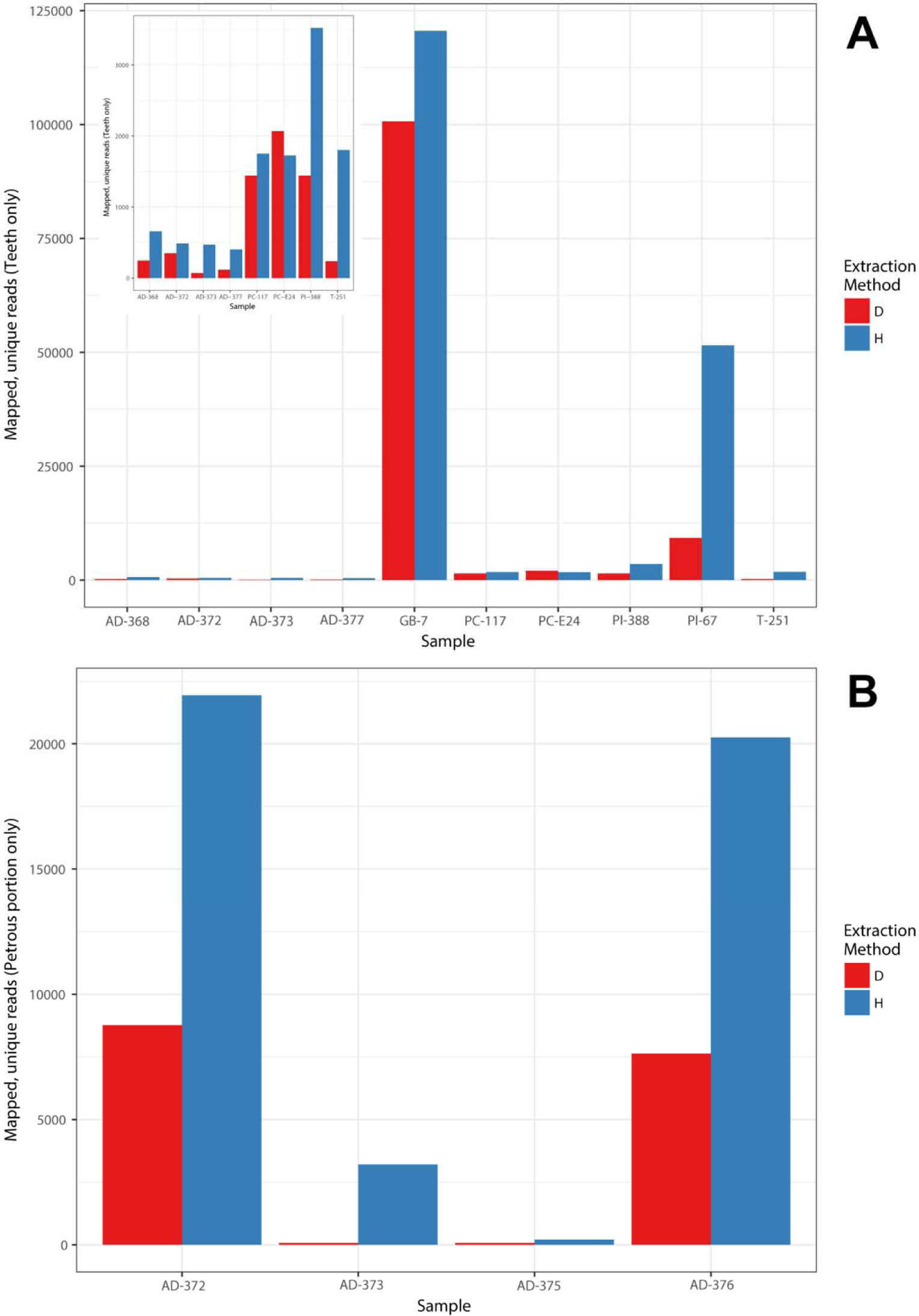
Number of mapped, unique reads per sample. (A) Tooth samples, inset zooms in to samples with less than 60,000 reads. (B) Petrous portion samples.

Sequence clonality (measured as fraction of the mapped reads that are duplicates) ranged between 0% to 14%, and tended to be higher for Method H libraries (Figure 2, Figure S5). Percent clonality was found to be significantly different between extraction methods for teeth, but not petrous portion samples (Table 2). The relationship between clonality and endogenous content is shown in Figure S6A. It was not linear or significant for either tissue (Teeth: Pearson’s *r* =−0.1265, t = −0.5409, df = 18, p=0.5952; Petrous portions: Pearson’s *r* =−0.2466, t = −0.6234, df = 6, p=0.5559).

**Figure 2.**
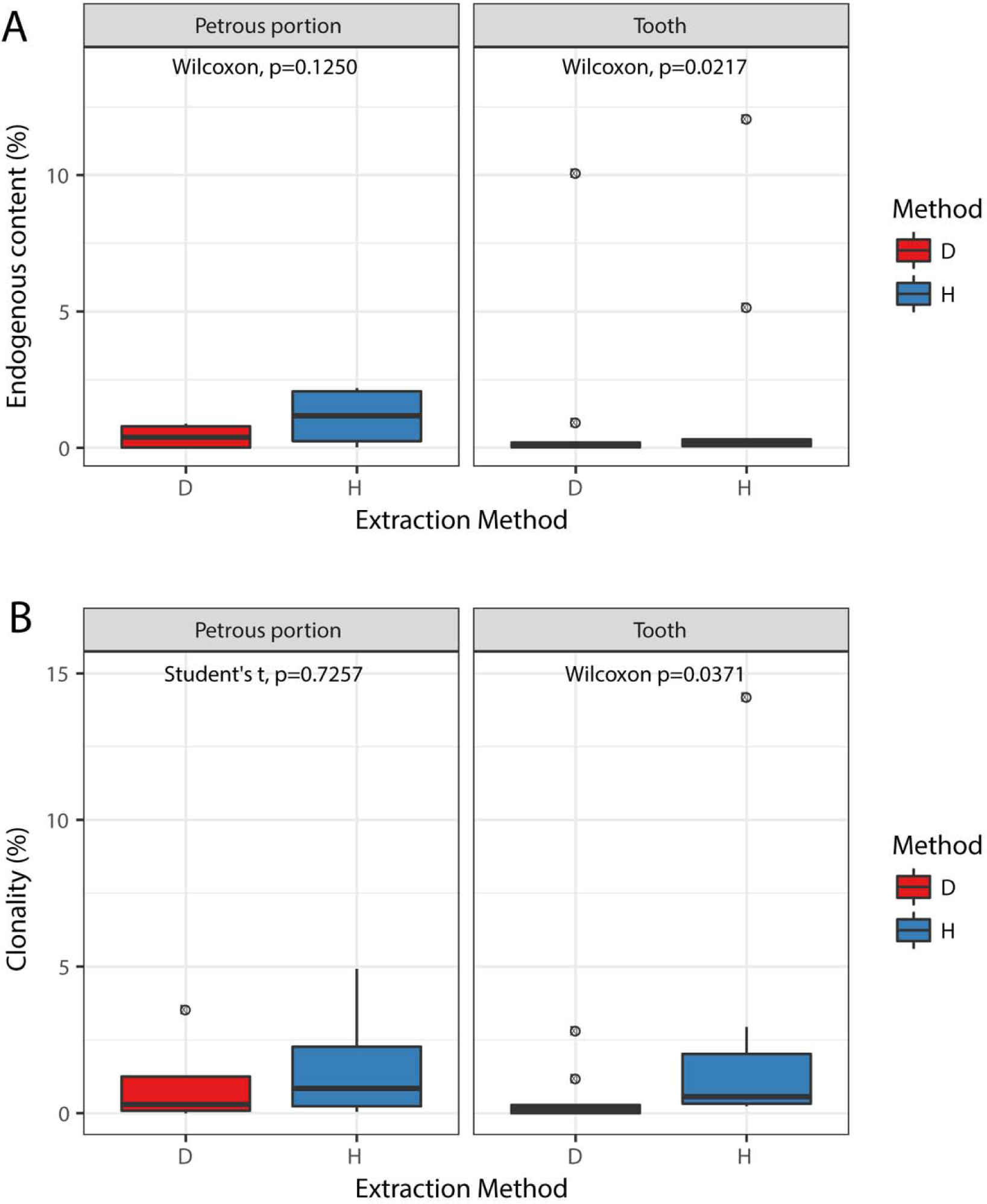
Boxplots comparing distributions of (A) percent endogenous content (calculated as number of mapped, unique reads divided over total down sampled reads) and (B) percent clonality (calculated as fraction of downsampled reads that are duplicates).

To examine this question further and test which method produced higher complexity libraries, we used the *c_curve* function in preseq to estimate the number of distinct reads recovered for each library. High complexity libraries have a large proportion of distinct reads that map to different parts of the reference genome. Therefore, more parts of the reference are covered with a single sequencing experiment. In contrast, low complexity libraries have a large proportion of distinct reads that map to the same sites and therefore may have a strong bias and high redundancy (Head et al., 2014). In this dataset, complexity was high regardless of extraction method. Method D libraries had a slightly higher mean proportion of distinct reads than Method H libraries, irrespective of tissue type, but this difference was not statistically significant (Figure S7). The relationship between complexity and endogenous content in the tested libraries is shown in Figure S6B. No significant correlation was observed between the two values (Teeth: Pearson’s *r* = 0.1288, t = 0.5514, df = 18, p=0.5881; Petrous portions: Pearson’s *r* = 0.3789, t = 1.003, df = 6, p=0.3546).

We used the *lc_extrap* function within preseq to predict the expected yield for a larger sequencing effort with the same libraries. This extrapolation analysis is highly sensitive to the amount of sequence data generated, and can give false estimates with low amounts of reads (Daley & Smith, 2013). Therefore, this analysis was only possible for sample-treatment combinations in the two libraries that yielded the highest number of reads: PI-67 and GB-7. Figure 3 demonstrates that, in both cases, libraries constructed with Method H extracts were predicted to yield a higher amount of complex DNA fragments with deeper sequencing (up to 600 million reads). In both cases, saturation of the complexity curve is reached earlier for libraries created from Method D extracts.

**Figure 3.**
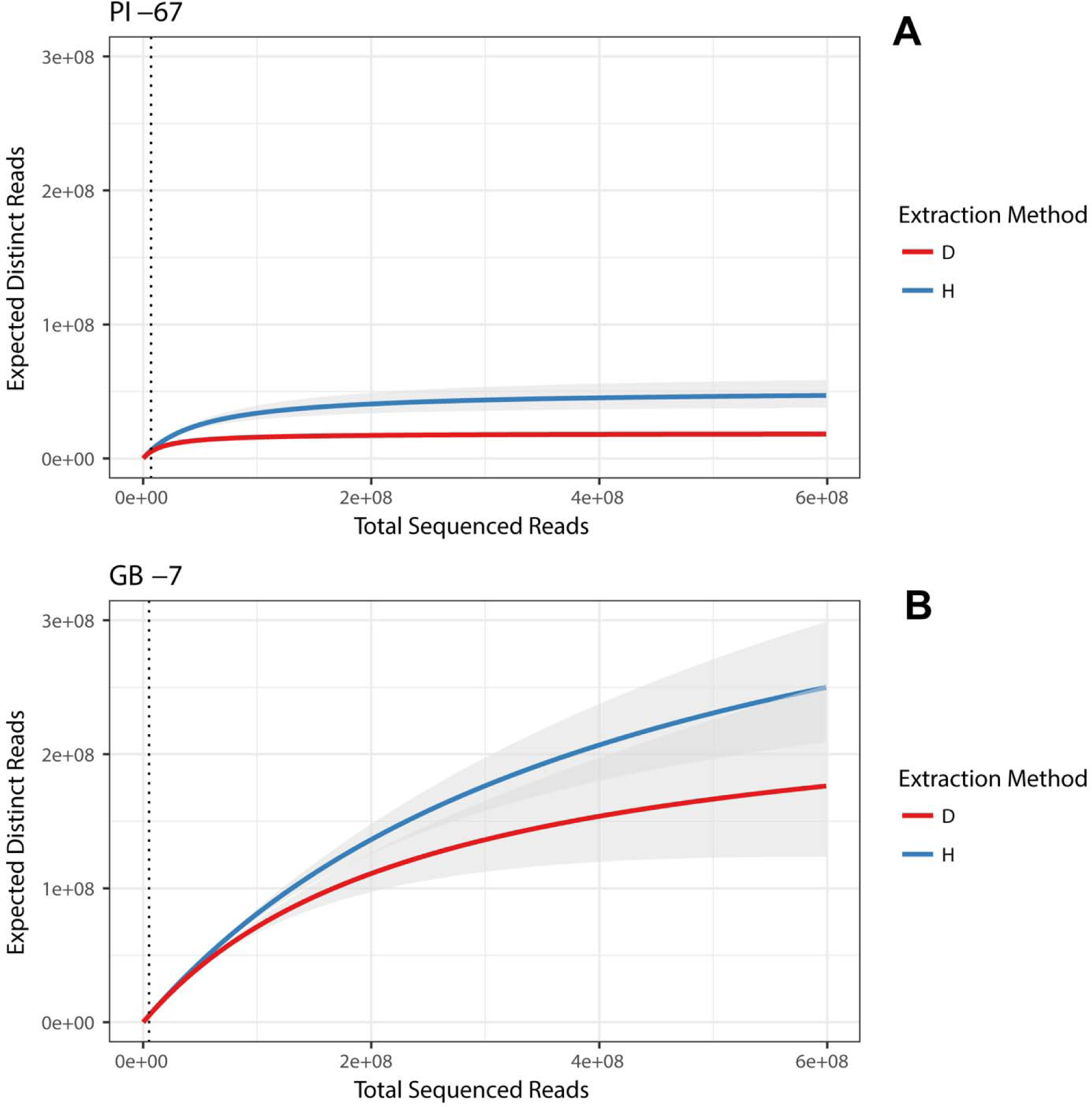
Extrapolation curves for shotgun library complexity estimation. Curves are shown for the two samples with highest number of reads: PI-67 and GB-7. Top inset shows zoomed in results for PI-67. Extrapolation curve and confidence intervals were estimated in preseq using default parameters and assuming a sequencing effort of 600 million reads. The dotted line denotes the number of reads randomly downsampled for each sample pair: 5.1 million reads for GB-7 and 6.8 million reads for PI-67.

### DNA fragment lengths and GC content

All samples, irrespective of extraction method had average DNA fragment lengths <100 bp. This small size is consistent with expectations for degraded remains (Briggs et al., 2007; Dabney et al., 2013b; Meyer et al., 2014) and similar to sizes obtained in previous aDNA research with tropical samples (Kehlmaier et al., 2017; Schroeder et al., 2015). Samples extracted with Method D yielded smaller average DNA fragment sizes (Tooth: 58.63 bp and Petrous portion: 53.65 bp) than those extracted with Method H (Tooth: 77.22 and Petrous portion: 72.07 bp) (Figure 4A). Overlaid plots showing the length distribution of sequence reads, both before and after mapping and filtering, demonstrate that Method H libraries had a higher proportion of larger fragments (Figure S8-10). Although this pattern of higher average fragment lengths in Method H libraries is evident in boxplots for both tooth and petrous portion samples, this difference was only found to be statistically significant in teeth (Figure 4A). We suspect this finding is influenced by low statistical power due to the smaller size of the petrous portion sample. No significant correlation was found between endogenous content and read length in either tissue type (Teeth: Pearson’s *r* = 0.3321 t = 1.4936, df = 18, p=0.1526; Petrous portions: Pearson’s *r* = 0.2647, t = 0.6724 df = 6, p=0.5263) (Figure S11A).

**Figure 4.**
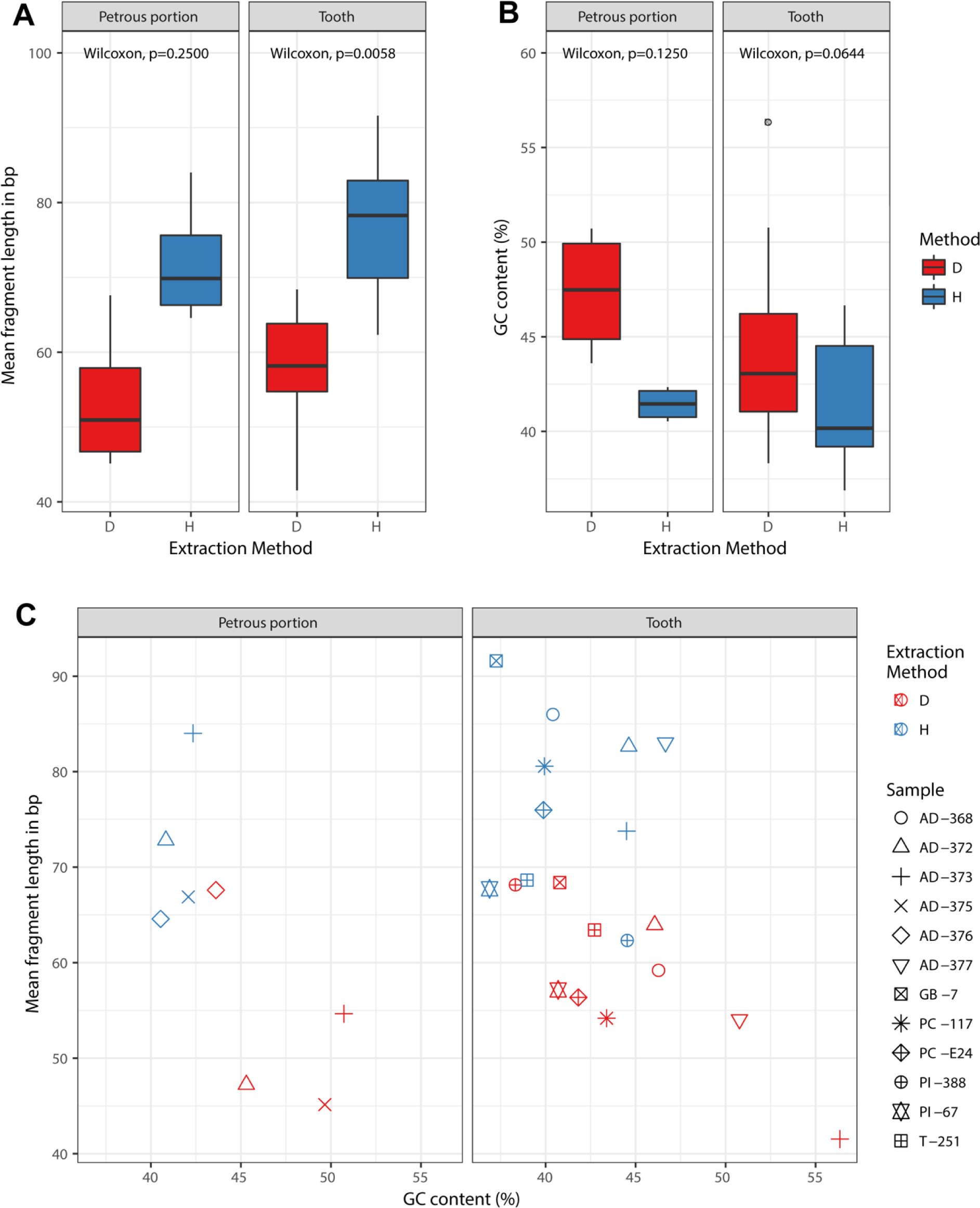
Fragment length and GC content. (A) Boxplots comparing distributions of DNA fragment lengths. (B) Boxplots comparing distributions of average %GC content. (C) Scatterplot of mean fragment length versus average %GC content.

Method D libraries had higher GC content irrespective of tissue, but this difference was not found to be statistically significant (Table 2, Figure 4B). A scatterplot of average DNA fragment lengths versus average percent GC content clearly distinguishes between samples generated with each method for both tissues (Figure 4C). A significant negative correlation was found between average DNA fragment length and GC content for all samples (Teeth: Pearson’s *r* = −0.5585, t = −2.8566, df = 18, p=0.01048; Petrous portions: Pearson’s *r* = −0.7181, t = −2.5278 df = 6, p=0.0448). No significant correlation was observed between percent GC and endogenous content (Teeth: Pearson’s *r* = −0.4092, t = −1.903 df = 18, p=0.0732; Petrous portions: Pearson’s *r* = −0.6578, t = −2.1395, df = 6, p=0.0762) (Figure S11B).

### DNA damage

Neither of the three DNA damage patterns examined (λ,δ_D_, δ_S_) differed significantly between samples extracted with either extraction method (Figure S12). Most samples had high probabilities (>0.70) of C to T and G to A misincorporations caused by DNA damage at the first and last position of each fragment (see Supplementary File 3 for damage plots). This is consistent with known damage patterns of authentic aDNA sequences (Briggs et al., 2007; Dabney et al., 2013b; Overballe-Petersen et al., 2012). Replicates PC-117-D (tooth), AD-373-H (petrous portion) and AD-377-H (tooth) had the lowest damage patterns (~0.50 probability of C to T or G to A misincorporations). Coverage of the autosomal and mitochondrial genomes were insufficient for contamination determination with Bayesian tools which require >3-5X minimum read depth for confident assessment (Fu et al., 2013; Racimo et al., 2016; Renaud et al., 2015). However visual examination of BAM files did not reveal patterns consistent with contamination, such as multiple populations of sequence reads or high mismatch rates.

## DISCUSSION

In this research, we explored the performance of two extraction protocols on poorly preserved tooth and petrous portion remains excavated from tropical contexts. Our experimental results suggest that both Method D and Method H successfully recovered degraded genetic material from tooth and petrous portion samples. No statistically significant difference was observed in raw DNA yields, library complexity or postmortem damage patterns in shotgun reads. The latter suggests that neither method is biased against recovery of degraded DNA fragments. This finding is consistent with results previously reported by Gamba et al. (2016), who found that ancient samples extracted with several silica-based extraction methods did not exhibit different postmortem damage patterns. Other studies have demonstrated that modifying digestion or pre-digestion wash steps also had negligible effects on DNA damage profiles (Boessenkool et al., 2016; Damgaard et al., 2015).

Significant differences between methods were observed in DNA fragment length, endogenous content, clonality and number of mapped, unique reads (post-quality filtering). DNA fragments recovered with Method H were, on average, 19 base pairs longer than those recovered with Method D, irrespective of tissue. These differences were visible in boxplots for all tissue types but were only significant for comparisons with tooth extracts. The small size of the petrous portion sample likely resulted in low statistical power to detect significant differences. Because of this, we refrain from extrapolating meaningful conclusions based on the petrous portion datasets and note that further research with more comprehensive samples of petrous portion tissue is needed to resolve this question. The following discussion focuses on statistically significant trends observed in tooth samples only.

Extraction Method D was designed for recovery of ultrashort DNA fragments (Dabney et al., 2013a). Given that Method H uses the same binding and purification steps implemented in Method D, short DNA fragment loss may have occurred during the pre-digestion EDTA wash or during the extended digestion and decalcification step. Warinner et al. (2014) did not observe reductions in raw DNA yields after pre-extraction washes of calculus samples with EDTA. Other studies conducted with tooth and bone tissue also found no significant differences in average DNA fragment lengths recovered after modifying extraction procedures with extended digestion steps or bleach-based decontamination washes (Damgaard et al., 2015; Gamba et al., 2016; Korlevic et al., 2015). But a more recent report by Glocke and Meyer (2017) found that EDTA interferes with recovery of short DNA fragments. In general, reports comparing aDNA extraction methods have suggested that digestion time may be a strong influence on recovery rates and characteristics of endogenous aDNA. For instance, Damgaard et al. (2015) observed diminished aDNA recovery with digestion steps longer than one hour. More recently, Boessenkool et al. (2016) found that mean aDNA fragment lengths were smaller and GC content was higher in bone extractions performed with short digestions.

Method H implements an initial wash of 0.5M EDTA solution followed by an extended two-part digestion step in which bone powder is first incubated for one hour in lysis buffer, and then further kept overnight at 37°C. It is possible that further optimization of the EDTA wash solution and of subsequent digestion conditions, such as temperature and incubation time, are needed to avoid loss of small DNA fragments. Future optimization efforts may also benefit from separate library preparation and sequencing of EDTA wash and pre-digestion fractions to identify where small DNA fragments are being lost in the extraction process.

We additionally found that libraries built from Method H extracts had higher endogenous content and more reads mapping to the reference after quality-filtering. But, on average, these samples also had higher clonality. In other words, Method H libraries had more PCR duplicates and sequence reads with the same starting and ending coordinates. However, extrapolation of predicted library complexity with the two best preserved samples (PI 67 and GB 7) indicated that Method H libraries would yield more unique DNA fragments upon repeated sequencing experiments. Complexity analyses are highly sensitive to the amount of sequence data generated. Low amounts of reads can lead to false estimates due to uncertainty of the extrapolation (Daley & Smith, 2013). Examination of the complexity curves for both methods demonstrates that despite the high clonality of Method H libraries, deeper sequencing would likely be most useful with Method H versus Method D extracts. This pattern may be due to the higher number of unique reads after quality filtering that were recovered with Method H in the two samples examined (Table S2).

Lastly, although not a significant trend, we observed that average GC content was at least three percentage points higher in paired tooth samples extracted with Method D versus Method H. But all samples, irrespective of extraction method, showed a decrease in GC content with larger average fragment size. This finding is consistent with previous research which has shown that differential DNA preservation can cause compositional bias towards higher GC content in ancient genomes composed of short DNA fragments (Briggs et al., 2007; Glocke & Meyer, 2017; Krause et al., 2010; Schuenemann et al., 2011). Higher GC content has also been correlated with lower contamination due to reduced presence of exogenous DNA (Racimo et al., 2016). As GC content can be strongly affected by amplification enzymes used in the library preparation process (Aird et al., 2011; Dabney & Meyer, 2012; Seguin-Orlando et al., 2015), all samples in this study were amplified with the same conditions so we consider this unlikely to explain the observed differences in base composition between extraction treatments. At a first glance, these results suggest that since Method D likely allowed for higher recovery rates of GC-rich DNA, it may be better suited for ancient tropical samples. However, we did not identify a significant correlation between percent GC and endogenous content (Figure S11B). High GC content can also lead to low sequence coverage in aDNA due to difficulty with mapping and alignment (Krause et al., 2010; Schuenemann et al., 2011). Although this problem may be alleviated somewhat by deep sequencing and high read depths, increased recovery of GC-rich DNA, may not necessarily lead to better results when read depth and coverage is inherently low, such as in poorly preserved tropical samples.

Several recent studies have found that the combination of extraction methods geared towards ultrashort DNA fragment recovery with single-stranded library preparation substantially increase endogenous aDNA yields (Barlow et al., 2016; Glocke & Meyer, 2017). While we are cognizant of these recent advances, in this work we have focused on libraries built with the more commonly used double stranded library protocol (Meyer & Kircher, 2010). Except for the historic chimpanzee sample, all archaeological remains examined in this study contained very low levels of endogenous DNA (<2%). This suggests that enrichment approaches are essential for increasing informative sequence content with poorly preserved tropical samples (Carpenter et al., 2013; Schroeder et al., 2015). Ongoing research in our laboratory has found that aDNA samples from Puerto Rico contain sufficient endogenous DNA for effective enrichment of complete mitochondrial DNA genomes (Nieves-Colon et al., 2016). Thus, here we follow recommendations by Wales et al. (2015) and focus on double-stranded library protocols which are better suited for studies geared towards enrichment of multi-copy organellar DNA. Additionally, previous reports have demonstrated that extremely short molecules obtained after single-stranded library preparation (Gansauge et al., 2017; Gansauge & Meyer, 2013; Glocke & Meyer, 2017), are often lost during target enrichment capture (Ávila-Arcos et al., 2015).

## CONCLUSIONS

This study has demonstrated that Method D and Method H were similarly efficient in recovering endogenous DNA in archaeological and historic skeletal samples from tropical contexts. However, significant differences exist in the composition of the recovered sequence data. Although libraries from Method H yielded more unique sequence reads and higher endogenous content, libraries built with this method also had higher clonality and yielded more PCR duplicates. In contrast, Method D recovered smaller aDNA fragments. Because of the exacerbated aDNA degradation that takes place in the tropics, we expect most informative sequence content to come from small DNA fragments in ancient remains (Allentoft et al., 2012; Hofreiter et al., 2015). Therefore, our findings suggest that, until further optimization of new protocols can take place, Method D continues to be the optimal choice for maximizing aDNA recovery in tropical tooth samples from ancient or historic contexts.

We also note that the insights derived from this work are restricted to tooth samples only. The smaller sample sizes obtained for petrous portion samples did not allow for conclusive statements regarding each method’s performance with this tissue type. Future efforts to develop methodologies tailored for tropical aDNA samples will benefit from increased sampling of suitable petrous portions and other tissues, such as dental calculus. A larger dataset shall further allow for finer sub settings of the data so that study results will control for differences between relatively well versus poorly preserved samples and/or for differences in site-specific aDNA preservation patterns.

## ACKNOWLEDGEMENTS

The authors would like to thank the research team at the Gombe National Park, Dr. Mike Wilson, Dr. Anne Pusey, and Dr. Ian Gilby for facilitating research with the Gombe chimpanzees. We especially thank Dr. Jane Goodall for ensuring the preservation of skeletal material and making them available for further study, and Dr. Rebecca Nockerts for preparing the samples. In addition, we also thank Dr. L. Antonio Curet and Dr. Tim Webster for helpful discussion, and the support staff and volunteers at the Ceremonial Center of Tibes Archaeological Park, and the Yaxuna archaeological site for invaluable logistics support. Lastly, we thank the three reviewers whose thoughtful comments and suggestions led to improvement of this paper.

Research with Puerto Rico samples was supported by National Science Foundation Grant BCS-061272 to W.J.P., National Science Foundation Doctoral Dissertation Improvement Award BCS-1622479 and Rust Family Foundation grant in Archaeology to M.N.C. Funding for work with Gombe samples was provided by the Leakey Foundation, ASU Strategic Initiative Funds, Office of the President of Arizona State University to A.C.S, and the Institute of Human Origins’ *DNA and Human Origins at ASU* project. We thank the Consejo de Arqueología of the Instituto Nacional de Antropología e Historia for granting the permits to conduct the research at Yaxuna, which was generously supported by the Fundación Roberto Hernández, Fundación Pedro y Elena Hernández, the Selz Foundation, and Jerry Murdoch. M.N.C wishes to further thank the Arizona State University School of Human Evolution and Social Change Dissertation Completion Fellowship program for providing financial assistance during the time this work was completed.

Author contributions were as follows: M.N.C. and A.C.S. designed the research, W.J.P, A.C., V.T. and T.S. provided samples and taphonomic information, M.N.C. and A.O. performed the experiments, M.N.C. analyzed the data, A.C.S and T.S. provided materials and resources, M.N.C. wrote the article, with input from co-authors. Genetic data are available in the NCBI Short Read Archive (SRA) under BioProject PRJNA398385.

